# Divalent cation and chloride ion sites of chicken acid sensing ion channel 1a elucidated by x-ray crystallography

**DOI:** 10.1101/379685

**Authors:** Nate Yoder, Eric Gouaux

## Abstract

Acid sensing ion channels (ASICs) are proton-gated ion channels that are members of the degenerin/epithelial sodium channel superfamily and are expressed throughout central and peripheral nervous systems. ASICs have been implicated in multiple physiological processes and are subject to numerous forms of endogenous and exogenous regulation that include modulation by Ca^2^+ and Cl^−^ ions. However, the mapping of ion binding sites as well as a structure-based understanding of the mechanisms underlying ionic modulation of ASICs have remained elusive. Here we present ion binding sites of chicken ASIC1a in resting and desensitized states at high and low pH, respectively, determined by anomalous diffraction x-ray crystallography. The acidic pocket serves as a nexus for divalent cation binding at both low and high pH, while we observe divalent cation binding within the central vestibule on the resting channel at high pH only. Moreover, neutralization of residues positioned to coordinate divalent cations via individual and combined Glu to Gln substitutions reduced, but did not extinguish, modulation of proton-dependent gating by Ca^2+^. Additionally, we demonstrate that anion binding at the canonical thumb domain site is state-dependent and present a previously undetected anion site at the mouth of the extracellular fenestrations on the resting channel. Our results map anion and cation sites on ASICs across multiple functional states, informing possible mechanisms of modulation and providing a blueprint for the design of therapeutics targeting ASICs.

## Introduction

Acid sensing ion channels (ASICs) are voltage-insensitive and proton-gated(1) members of the epithelial sodium channel/degenerin (ENaC/DEG) superfamily of ion channels(2, 3) that assemble as homo- or heterotrimeric sodium-selective ion channels(4) and are expressed throughout vertebrate central and peripheral nervous systems. ASICs exhibit a simple three-state gating scheme, populating a non-conducting resting state at high pH, opening upon exposure to protons and quickly desensitizing(5), recovering proton-sensitivity only upon return to high pH. ASICs gate rapidly, fully activating in milliseconds and undergoing complete desensitization in hundreds of milliseconds(6, 7). The homotrimeric splice variant ASIC1a is highly enriched in the central nervous system (CNS) and participates in numerous physiological processes including learning and memory(8) and nociception(9). Furthermore, ASIC1a is moderately permeable to Ca^2+^ and has been implicated in various forms of acidosis-induced neuronal injury and neurological disorders(10, 11).

ASIC1a activity is modulated by endogenous divalent cations including Zn^2+^, Mg^2+^, and Ca^2+^(5, 12–14) and the modification of ASIC1a gating by extracellular Ca^2+^ has been an area of active investigation. Previous studies of homotrimeric ASIC1a and ASIC3 channels have proposed both allosteric(5) and pore blockade(13, 15, 16) mechanisms and suggested correspondingly distinct binding sites (13). Furthermore, residues corresponding to Glu 426 and Asp 433 of cASIC1a are located at the mouth of the extracellular fenestration and within the pore, respectively, and are critical to Ca^2+^-dependent block(13). However, ambiguous results and a lack of defined Ca^2+^ binding sites have fallen short of a comprehensive mechanism for gating modification of ASIC1a channels by Ca^2+^.

Extracellular Cl^−^ ions modulate a variety of ASIC1a characteristics including proton-dependent gating, desensitization kinetics and tachyphylaxis(17). In x-ray structures of ASIC1a channels in open and desensitized states, a bound Cl^−^ is buried within the thumb domain near helices α4 and α5(4, 18, 19). At low pH, Cl^−^ ions are positioned at a subunit-subunit interface coordinated by Arg and Glu residues on α4 of the thumb domain and by a Lys residue on the palm domain of a neighboring subunit. Residues involved in Cl^−^ binding are highly conserved amongst ASICs, and modulation by extracellular Cl^−^ has also been observed in ENaC channels at a likely similar inter-subunit binding site(20, 21), demonstrating the importance of extracellular Cl^−^ to the ENaC/DEG superfamily of ion channels.

Here, we determine binding sites for anions and divalent cations on resting and desensitized ASIC1a channels at high and low pH, respectively. Our results map a complex network of divalent cation sites on ASICs within domains closely involved in proton-dependent gating and demonstrate state-dependence for both anion and cation binding. These data present a structural framework for understanding the interplay between gating and ion binding in ASICs and provide a template for the development of ASIC1a-specific modulatory agents.

## Materials And Methods

### Receptor construct, expression and purification

The Δ25 and Δ13 crystallization constructs have 24 or 13 residues removed from the amino-terminus, respectively, and 64 residues removed from the carboxy-terminus(19, 22, 23). Recombinant protein was expressed in HEK293S GnTI^−^ cells(24) as previously described(23). In brief, HEK293S GnTI^−^ cells grown in suspension were infected with BacMam virus(25) and collected by centrifugation after 48 hours of culture. Cell pellets were resuspended in tris buffered saline (TBS; 150 mM NaCl, 20 mM Tris pH 8.0, 1 mM phenylmethylsulfonyl fluoride, 0.05 mg ml^-1^ aprotinin, 2 μg ml^-1^ pepstatin A, and 2 μg ml^-1^ leupeptin), disrupted by sonication, and membrane fractions were isolated by ultracentrifugation.

Membrane pellets were solubilized in TBS with 40 mM *n*-dodecyl β-D-maltoside (DDM) and clarified by ultracentrifugation. Solubilized membranes were incubated with Co^2+^ affinity resin and protein was eluted with buffer containing 300 mM NaCl, 20 mM Tris pH 8.0, 1 mM DDM, and 250 mM imidazole. Protein purification buffers for Δ13 were identical to those used for Δ25 but contained 150 mM NaCl. The histidine-tagged enhanced green fluorescent protein (EGFP) tag was cleaved with thrombin digestion and the protein was further purified by size-exclusion chromatography using a mobile phase containing 300 mM NaCl, 20 mM Tris pH 8.0, 2 mM n-decyl β-D-thiomaltopyranoside (C10ThioM), 1 mM dithiothreitol (DTT), 0.2 mM cholesteryl hemisuccinate (CHS) and 5 mM BaCl_2_. Δ13 protein was further purified by size-exclusion chromatography (SEC) using a mobile phase containing 150 mM NaCl, 20 mM Tris pH 8.0, 1 mM DDM, 1 mM DTT, and 0.2 mM CHS. Peak fractions were collected and concentrated to ~2-3 mg ml^-1^. The source for all cell lines was ATCC.

### Crystallization

Crystallization of purified Δ25 protein was accomplished as previously described(23). Purified Δ13 protein was used for crystallization trials immediately following SEC and was not subject to any additional treatment. The Δ13 crystals were obtained at 20°C by way of the hanging drop vapor diffusion method. Reservoir solution contained 100 mM HEPES pH 6.9, 150 mM sodium formate and 18% (w/v) PEG 3,350. Drops were composed of 1:1, 1:2, and 2:1 protein to reservoir ratios, respectively. Crystals typically appeared within 2 weeks. Crystals were cryoprotected with 30% (v/v) glycerol, in the protein-containing drop, before flash cooling in liquid nitrogen.

### Anomalous scattering experiments

To locate Ba^2+^ sites on Δ25 channels, crystals were grown using a reservoir solution composed of 150 mM NaCl, 100 mM Tris pH 8.5, 20 mM BaCl_2_, and 29% PEG 400 (v/v). The Δ25 crystals were soaked in solution containing 150 mM NaCl, 100 mM Tris pH 8.5, 50 mM BaCl_2_, 36% PEG 400 (v/v), 2 mM C10ThioM and 0.2 mM CHS for 5 minutes prior to freezing in liquid nitrogen. To locate Cl^−^ sites on Δ25 channels, we exploited the anomalous scattering from bromide and grew crystals of Δ25 using a reservoir solution composed of 150 mM NaCl, 100 mM Tris pH 8.5, 5 mM BaCl_2_ and 33% PEG 400 (v/v). Crystals were soaked in solution containing 150 mM NaBr, 100 mM Tris pH 8.5, 5 mM BaCl_2_, 36% PEG 400 (v/v), 2 mM C10ThioM and 0.2 mM CHS for 2 minutes prior to freezing in liquid nitrogen. To locate Ba^2+^ sites on Δ13 channels, crystals were soaked in solution containing 150 mM sodium formate, 100 mM HEPES pH 6.9, 50 mM BaCl_2_, 30% (v/v) glycerol, 1 mM DDM and 0.2 mM CHS for 5 minutes prior to freezing in liquid nitrogen.

### Structure Determination

X-ray diffraction data sets were collected at the Advanced Light Source (ALS) beamline 5.0.2 and at the Advanced Photon Source (APS) beamline 24-ID-C. For anomalous diffraction experiments, diffraction data from crystals soaked in solutions containing 150 mM NaBr and belonging to the P2_1_2_1_2_1_ space group were measured using an x-ray beam tuned to 13,490 eV at APS beamline 24-ID-C. Crystals soaked in solutions containing 50 mM BaCl_2_ belonged to the P2_1_2_1_2_1_ space group (Δ25) and H3 (Δ13) and diffraction data were measured using an x-ray beam tuned to 6,400 eV at ALS beamline 5.0.2. Diffraction for both BaCl_2_ and NaBr-soaked crystals was measured to ~ 3.7-4.0 Å resolution.

Diffraction data were indexed, integrated, and scaled using XDS and XSCALE(26) software and structures were solved by molecular replacement using the PHASER program(27). To identify anomalous difference peaks in Ba^2+^-soaked and NaBr-soaked Δ25 crystals, coordinates from the Δ25-Ba^2+^ structure(23) (pdb 5WKU) were used as a search probe for molecular replacement in PHASER and anomalous difference Fourier maps were generated in Phenix(28). To identify anomalous difference peaks in Ba^2+^-soaked Δ13 crystals, coordinates from the desensitized state structure(18) (pdb 4NYK) were used as a search probe for molecular replacement. To generate anomalous difference maps, diffraction data was truncated at 5.0 Å for Ba^2+^ and 6.0 Å for Br^−^. Anomalous signals in the difference Fourier maps for Δ25 channels were enhanced by real-space averaging around the molecular 3-fold axis using Coot(29). Anomalous difference peaks were inspected and used to assign Ca^2+^ binding sites in the Δ25-Ca^2+^ structure(23) (pdb 5WKV). Only peaks stronger than 6 σ were considered for Ba^2+^ peaks and greater than 5 σ for Br^−^ peaks after non-crystallographic symmetry averaging, in the corresponding anomalous difference electron density maps. Accession codes 5WKX (Supporting information figure 1), 5WKY (Supporting information figure 2), and 6CMC (Supporting information figure 3) correspond to Ba^2+^ sites on Δ25, Br^−^ sites on Δ25 and Ba^2+^ sites on Δ13, respectively.

### Patch clamp recordings

Whole-cell patch clamp recordings of CHO-K1 cells expressing recombinant protein were performed as previously described(23). The Axopatch 200B amplifier and pClamp 10 software were used for data acquisition and trace analysis and only single recordings were taken from individual cells. In high Ca^2+^ experiments, both low pH and conditioning solutions contained 10 mM Ca^2+^. The source for all cell lines was ATCC.

## Results

### Crystallization of ASIC1a in resting and desensitized states

The Δ25 and Δ13 crystallization constructs span residues 25-463 and 14-463 of chicken ASIC1, a homolog of human ASIC1a, respectively, and maintain proton-dependent gating characteristics(19, 22, 23). Crystals of resting Δ25 channels belong to the P2_1_2_1_2_1_ space group and were obtained at high pH and in the presence of Ba^2+^. Crystals of desensitized Δ13 channels belong to the H3 space group and were obtained at low pH. All x-ray structures were determined by molecular replacement and built via iterative rounds of manual model building and refinement. All ion sites were confirmed via anomalous diffraction experiments using synchrotron radiation tuned to 6,400 eV for Ba^2+^ and 13,490 eV for Br^−^ (Table 1).

**Table 1:** Crystallographic and structure refinement statistics.

| | $\Delta 25$-$Ba^{2+}$ Resting<br>$Ba^{2+}$ - soaked<br>(5WKX) | $\Delta 13$-Desensitized<br>$Ba^{2+}$ - soaked<br>(6CMC) | $\Delta 25$-$Ba^{2+}$ Resting<br>$Br^-$ - soaked<br>(5WKY) |
| --- | --- | --- | --- |
| **Data collection** | ALS 502 | ALS 502 | APS 24ID-C |
| Space group | $P2_12_12_1$ | H3 | $P2_12_12_1$ |
| Cell dimensions<br>a, b, c (Å) | 108.0 126.3 156.3 | 133.9 133.9 122.7 | 109.6 134.5 159.1 |
| Cell angles<br>$\alpha, \beta, \gamma$ (°) | 90.0, 90.0, 90.0 | 90.0, 90.0, 120.0 | 90.0, 90.0, 90.0 |
| Wavelength (Å) | 1.94 | 1.94 | 0.9191 |
| Resolution (Å)* | 50 – 4.0 | 42.14 – 3.7 | 45 – 4.00 |
| Completeness* | 99.5 (97.1) | 99.61 (98.8) | 99.9 (99.4) |
| Multiplicity* | 4.1 (5.1) | 3.6 (3.5) | 7 (6.2) |
| $I/\sigma I$* | 8.82 (0.81) | 6.78 (0.98) | 9.46 (0.77) |
| $R_{meas}$ (%)* | 11.8 (326.9) | 15.2 (135.7) | 13.2 (206.6) |
| $CC_{1/2}$ (%)* | 99.9 (35.7) | 99.7 (31.6) | 100 (53.4) |
| **Refinement** |  |  |  |
| Resolution (Å) | 49.7 – 4.0 | 42.1 – 3.7 | 43.2 – 4.0 |
| No. of reflections | 17205 | 8949 | 20402 |
| $R_{work}/R_{free}$ | 30.2/31.3 | 28.4/30.0 | 28.1/29.9 |
| **Average B-factor (Å<sup>2</sup>)** |  |  |  |
| Protein | 142.0 | 133.0 | 133.0 |
| Ligand | 190.8 | 119.9 | 181.6 |
| **R.m.s. deviations** |  |  |  |
| Bond lengths (Å) | 0.004 | 0.004 | 0.004 |
| Bond angles (°) | 0.705 | 0.98 | 0.706 |
| **Ramachandran plot** |  |  |  |
| Favored (%) | 98.31 | 95.39 | 98.31 |
| Allowed (%) | 1.69 | 4.61 | 1.69 |
| Disallowed (%) | 0 | 0 | 0 |
| Rotamer outliers (%) | 1.24 | 1.78 | 1.24 |

\*Highest resolution shell in parentheses.
5% of reflections were used for calculation of $R_{free}$ .

### Cation binding sites on the resting channel at high pH

Gating of chicken ASIC1a channels is modulated by Ca^2+^, manifested as an acidic shift in pH50 with increasing concentrations of extracellular Ca^2+^ (Figure 1A). Despite extensive scrutiny, proposed mechanisms of gating modulation by Ca^2+^ remain uncertain. Inspection of the Fo-Fc density maps from the x-ray structure of a resting ASIC1a channel at high pH and in the presence of Ca^2+^(23) revealed strong positive difference density positioned within the electrostatically negative acidic pocket and central vestibule, regions of the channel involved in proton-dependent gating(30). In an effort to unambiguously assign binding sites for divalent cations on the resting channel structure, Δ25 crystals grown at high pH were soaked in Ba^2+^, whose effect on ASIC1a channels mimics that of Ca^2+^(12). Inspection of anomalous difference electron density maps subjected to real space threefold averaging revealed nine Ba^2+^ sites within the acidic pocket and the central vestibule (Figure 1B), in general agreement with electron density maps from the Δ25-Ca^2+^ x-ray structure. However, given the resolution limitations we are unable to define the protein atoms directly coordinating the Ba^2+^ ions and we therefore simply describe the protein residues that surround the ion sites.

**Figure 1.**
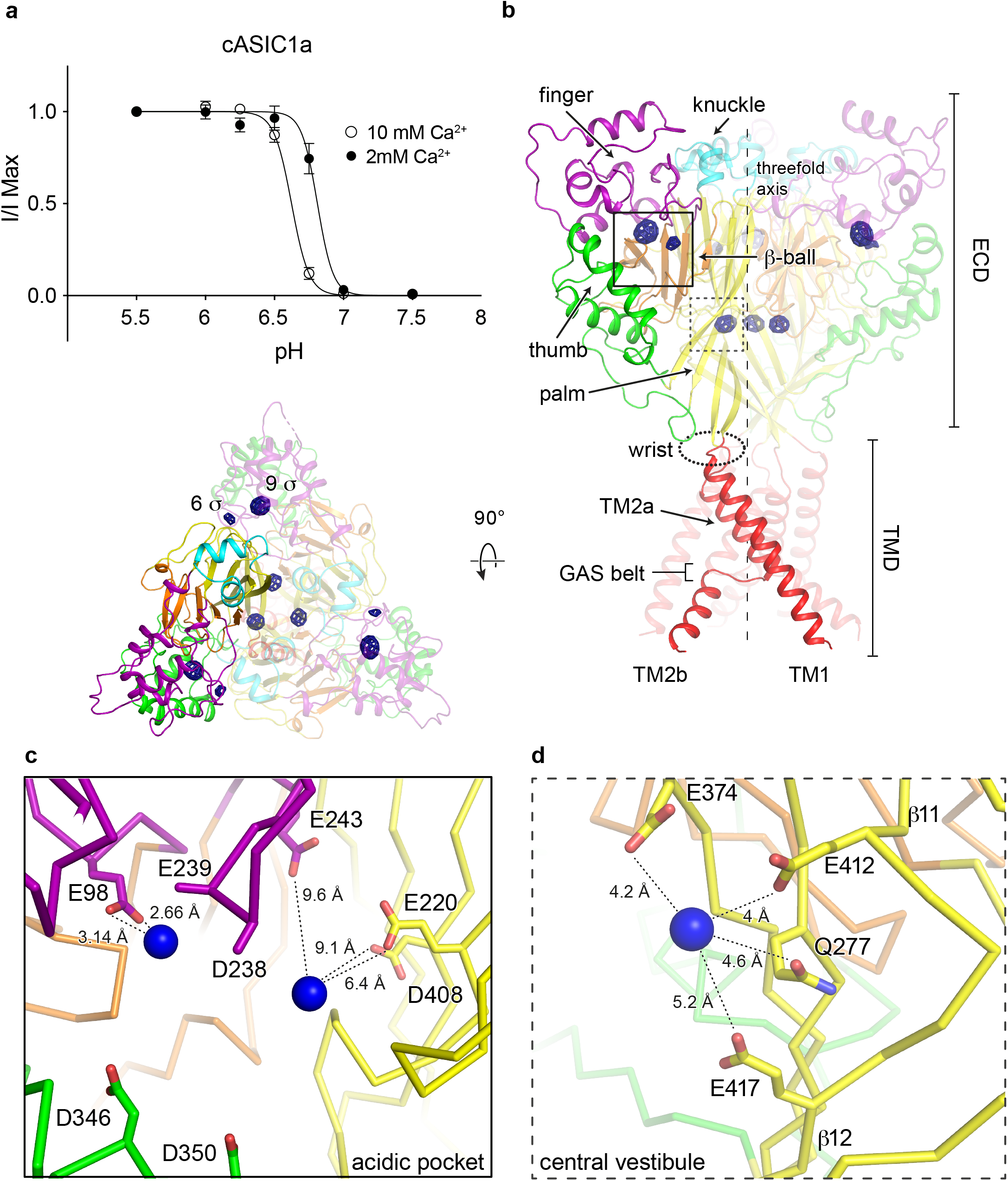
Binding sites for divalent cations on a resting chicken ASIC1a channel at high pH. **a,** Proton dose-response curves for cASIC1a channels in 2 or 10 mM external Ca^2+^. Error bars represent SEM, n = 4-8 cells. Data were collected in whole-cell patch clamp configuration from adherent CHO-K1 cells transfected with cDNA for cASIC1a channels. **b-d,** Ba^2+^ anomalous difference peaks (blue mesh) contoured at 5.5 σ and mapped on a resting channel at high pH (**b**, channel colored by domain) with detail views of the acidic pocket (**c**) and central vestibule (**d**) binding sites. Blue spheres represent Ba^2+^.

The acidic pocket, an electrostatically negative cavern which harbors binding sites for toxins (19, 22, 31) and putative proton sites, and undergoes a large conformational change during channel activation, contains two anomalous difference peaks for Ba^2+^ on the resting channel at high pH. The 9 and 6 σ anomalous difference peaks for Ba^2+^ are positioned ~ 8 Å apart at opposite corners of the acidic pocket, which is in an expanded conformation at high pH (Figure 1C). Electron density is poor within the solvent-exposed cavern, restricting modeling of some sidechains to α-carbon atoms and precluding a comprehensive picture of ion coordination. Despite this limitation, the position of the 9 σ anomalous difference peak deep within the acidic pocket suggests possible coordination of Ba^2+^ by acidic residues Glu 98 and Glu 239 of the channel’s finger domain. The weaker, 6 σ Ba^2+^ peak is situated at a subunit-subunit interface within the acidic pocket near highly conserved residues Glu 243 of the finger domain and both Glu 220 and Asp 408 from the palm domain of a neighboring subunit.

The central vestibule, buried within the channel core and situated along the threefold axis of resting ASIC1a channels, harbors a trio of threefold symmetric 8 σ anomalous difference peaks for Ba^2+^. These Ba^2+^ ions are situated at the nexus of upper and lower palm domains ~ 5 Å from Glu 412, Glu 417, Glu 374 and Gln 277 residues and immediately off of β11-β12 linkers (Figure 1D).

Recently published x-ray structures of chicken ASIC1a in a resting state at high pH highlighted the structural transitions underlying channel activation and desensitization(23). During the rapid transition from a non-conducting, resting state at high pH to the open state at low pH, the electrostatically negative acidic pocket collapses as helices α4 and α5 swing inwards towards the acidic loop of the finger domain (Figure 2A-B). Collapse of the acidic pocket fully occludes the 9 σ peak positioned off of Glu 98 and Glu 239, suggesting that occupancy of this site may be incompatible with a fully activated channel and may therefore provide a mechanism for gating modification. In contrast, the 6 σ peak positioned near the carboxy-terminal end of α5 remains solvent-exposed, indicating that occupancy of this site may not contribute to gating modification by divalent cations (Figure 2C). These results indicate a degree of state-dependence with respect to divalent cation binding at the acidic pocket and suggest that acidic residues surrounding the 9 σ anomalous peak, including Glu 98 and Glu 239, are likely relevant to cation binding and thus to divalent cation modulation of ion channel gating.

**Figure 2.**
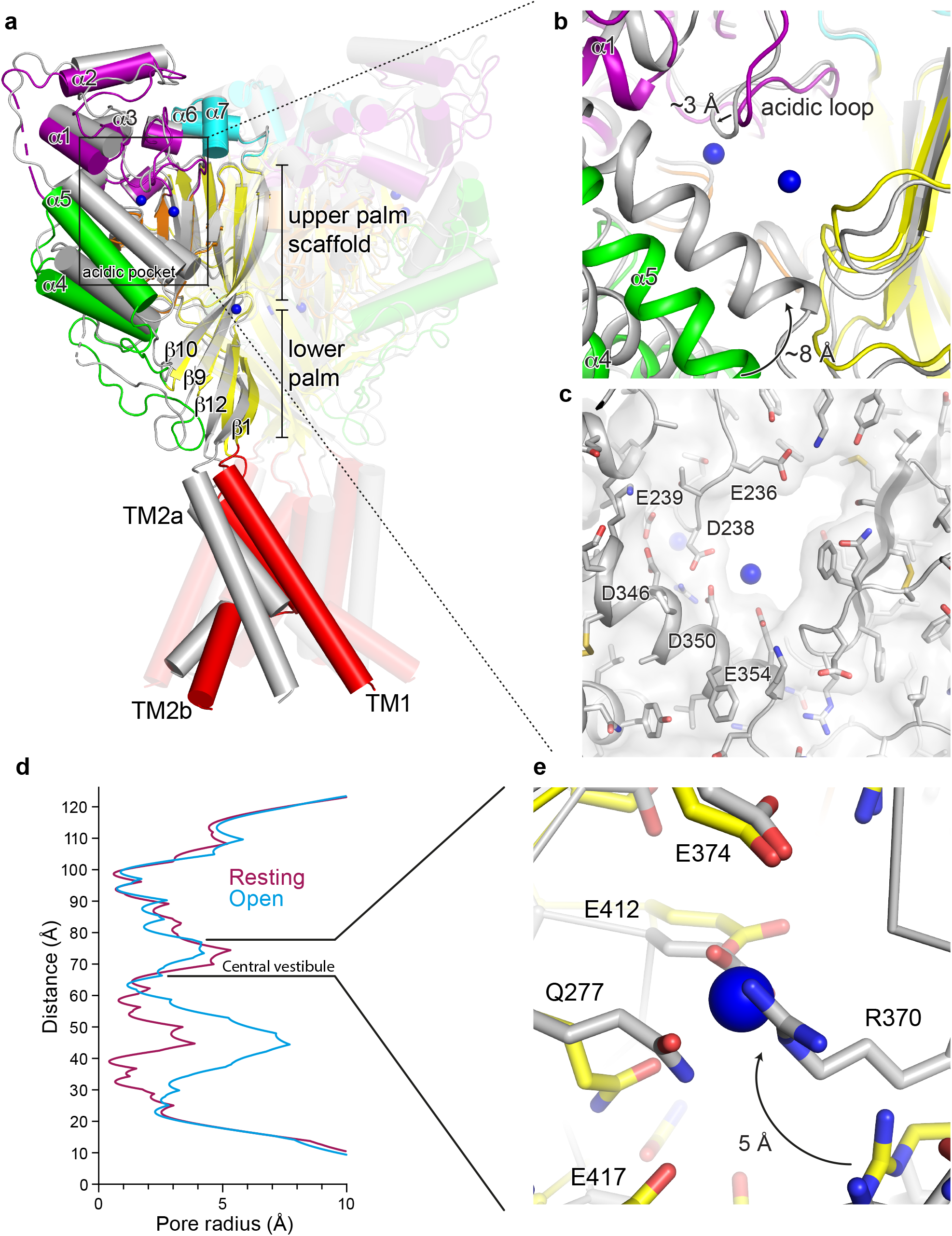
Activation initiates changes at binding sites for divalent cations. **a-b,** Superposition of resting and open (pdb 4NTW, grey) channels (**a**) with detail view of the acidic pocket (**b**). Blue spheres represent Ba^2+^ sites of the resting channel. **c,** Surface representation of the collapsed acidic pocket of the open channel (pdb 4NTW, grey). Blue spheres represent Ba^2+^ sites of the resting channel. **d,** Pore profiles for resting (pdb 5WKU) and open (pdb 4NTW) channels. **e,** Detail view of the central vestibule of superposed resting and open (pdb 4NTW, grey) channels. Blue spheres represent Ba^2+^ sites of the resting channel.

Structural changes during activation take place throughout the ECD, resulting in a slight contraction of the central vestibule as the lower palm domain flexes towards the membrane (Figure 2A, 2D). Consequently, Arg 370 within the central vestibule pivots towards the palm domain of a neighboring subunit upon exposure to low pH. Activation-induced reorientation of Arg 370 positions its guanidine group into a direct clash with the Ba^2+^ site on the resting channel at high pH (Figure 2E), rendering the open channel’s central vestibule not permissive to Ba^2+^ binding at the same site as in the resting channel. Because we do not have structural information on an open channel in complex with divalent cations, we cannot rule out the possibility of a unique binding site within the central vestibule that is only occupied in the open channel state.

These results suggest that binding sites for divalent cations within the central vestibule are state-dependent and may play a previously unanticipated role in the modification of ASIC gating by divalent cations. Moreover, the location of Ba^2+^ within the central vestibule of the resting channel overlaps that of the putative binding site for quercetin, an inhibitor of ASIC1a, ASIC2a, and ASIC3 homomers(32), suggesting that the central vestibule may serve as a common site for modulatory agents.

### Cation binding sites on a desensitized channel at low pH

To better explore the state-dependence of divalent cation binding to ASICs, we soaked crystals of the Δ13 construct of chicken ASIC1a grown at low pH in Ba^2+^, following a protocol identical to that used for Δ25 crystals grown at high pH. At low pH the Δ13 construct adopts the canonical desensitized channel structure(18), exhibiting a collapsed acidic pocket and closed gate. Inspection of anomalous difference electron density maps revealed a strong, ellipsoidal density for Ba^2+^ at the acidic pocket, indicating the presence of at least one bound ion (Figure 3A). Similar to the 6 σ site within the resting channel’s acidic pocket, the Ba^2+^ site of the desensitized channel is positioned near the carboxy-terminus of α5 at the mouth of the pocket. The proximity of numerous acidic residues indicates possible coordination of Ba^2+^ by Asp 238, Asp 350 and Glu 354, all of which are located between 3.4 and 4.3 Å from the Ba^2+^ site (Figure 3B). Notably, divalent cation sites within the resting channel’s central vestibule at high pH are absent on the desensitized channel at low pH. Moreover, despite a slight contraction at the central vestibule of the desensitized channel (Figure 3C), the conformation of the single Gln and three Glu residues that comprise the Ba^2+^ binding site on the resting channel site (Figure 3D) are remarkably similar in both resting and desensitized states. As such, the absence of an anomalous signal for Ba^2+^ for the desensitized channel, in addition to the presence of titratable resides lining the binding site, suggest that one or more of the aforementioned glutamate is protonated at low pH, thus disfavoring the binding of divalent cations.

**Figure 3.**
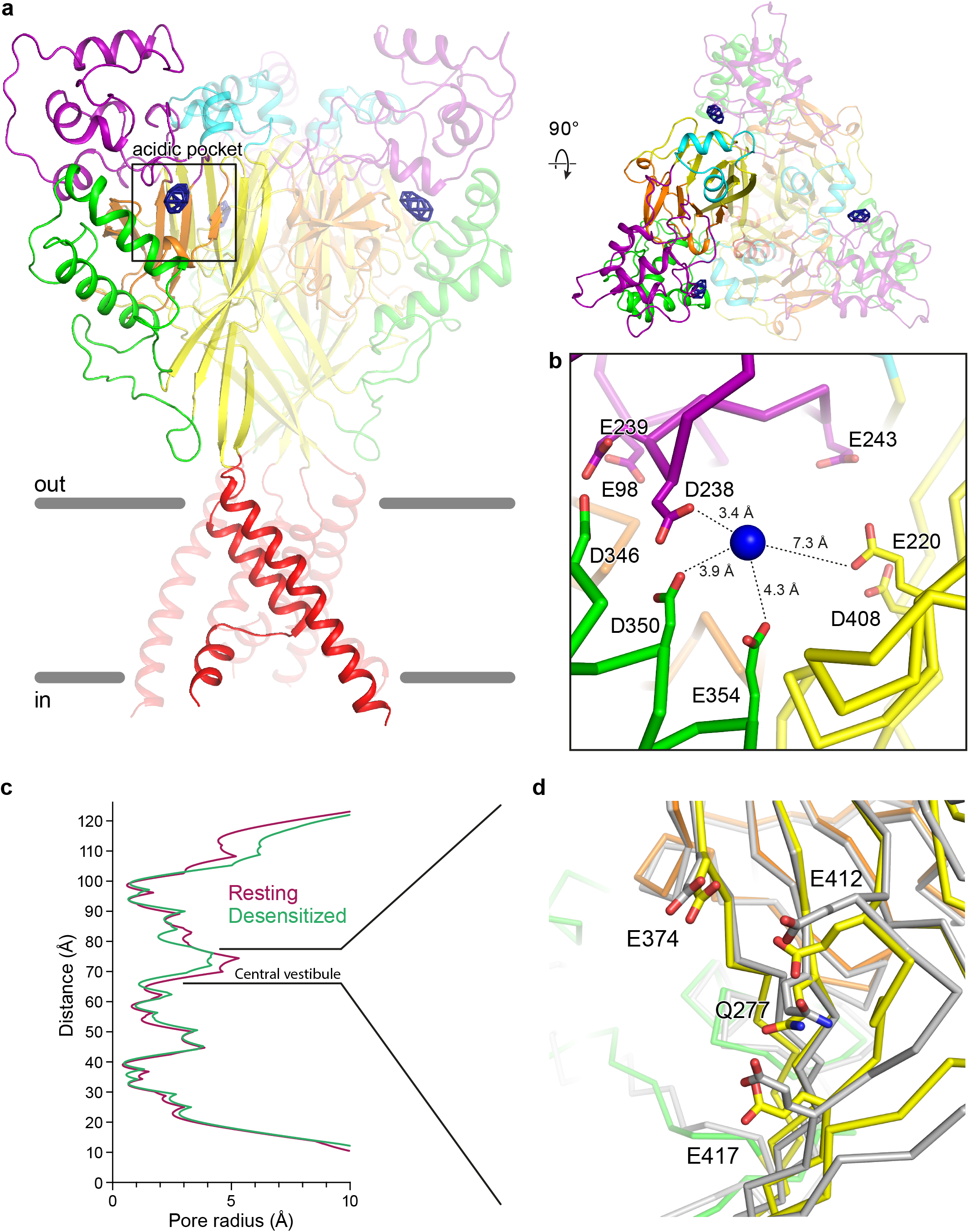
Binding sites for divalent cations on a desensitized chicken ASIC1a channel at low pH. **a,** Ba^2+^ anomalous difference peaks (blue mesh) contoured at 6.5 σ and mapped on a desensitized channel at high pH (channel colored by domain). **b,** Detail view of the Ba^2+^ binding site within the acidic pocket. Blue spheres represent Ba^2+^ sites of the desensitized channel. **c,** Pore profiles for resting (pdb 5WKU) and desensitized channels. **d,** Detail view of the central vestibule of superposed desensitized and resting (pdb 5WKU, grey) channels.

### Contribution of acidic residues to channel modulation

Inspection of electron density maps for the Δ25-Ca^2+^ x-ray structure (pdb 5WKV)(23), indicate that putative Ca^2+^ sites are in good agreement with anomalous difference peaks for Ba^2+^ on the resting channel at high pH. Within the acidic pocket and central vestibule of the Δ25-Ca^2+^ structure, Ca^2+^ ions are positioned off of Glu 98, Glu 220 and Glu 374 residues(23). To illuminate their contribution to channel modulation, we neutralized these three acidic residues via single E98Q, E220Q and E374Q as well as triple E98Q/E220Q/E374Q substitutions. In all mutant channels, increasing extracellular Ca^2+^ from 2 to 10 mM shifted the pH50 to more acidic values, though with less statistical significance than observed in cASIC1a channels (Figure 4A-B). The effect on modulation of gating by Ca^2+^ was least pronounced in the E220Q mutant, in good agreement with the observation from anomalous diffraction experiments that the divalent cation site within the acidic pocket near Glu 220 is occupied in both resting and desensitized states. Moreover, in contrast to all other mutant channels, neutralization of Glu 220 did not have a meaningful effect on pH50 at 2 mM Ca^2+^ when compared to cASIC1a (Figure 4A-B).

**Figure 4.**
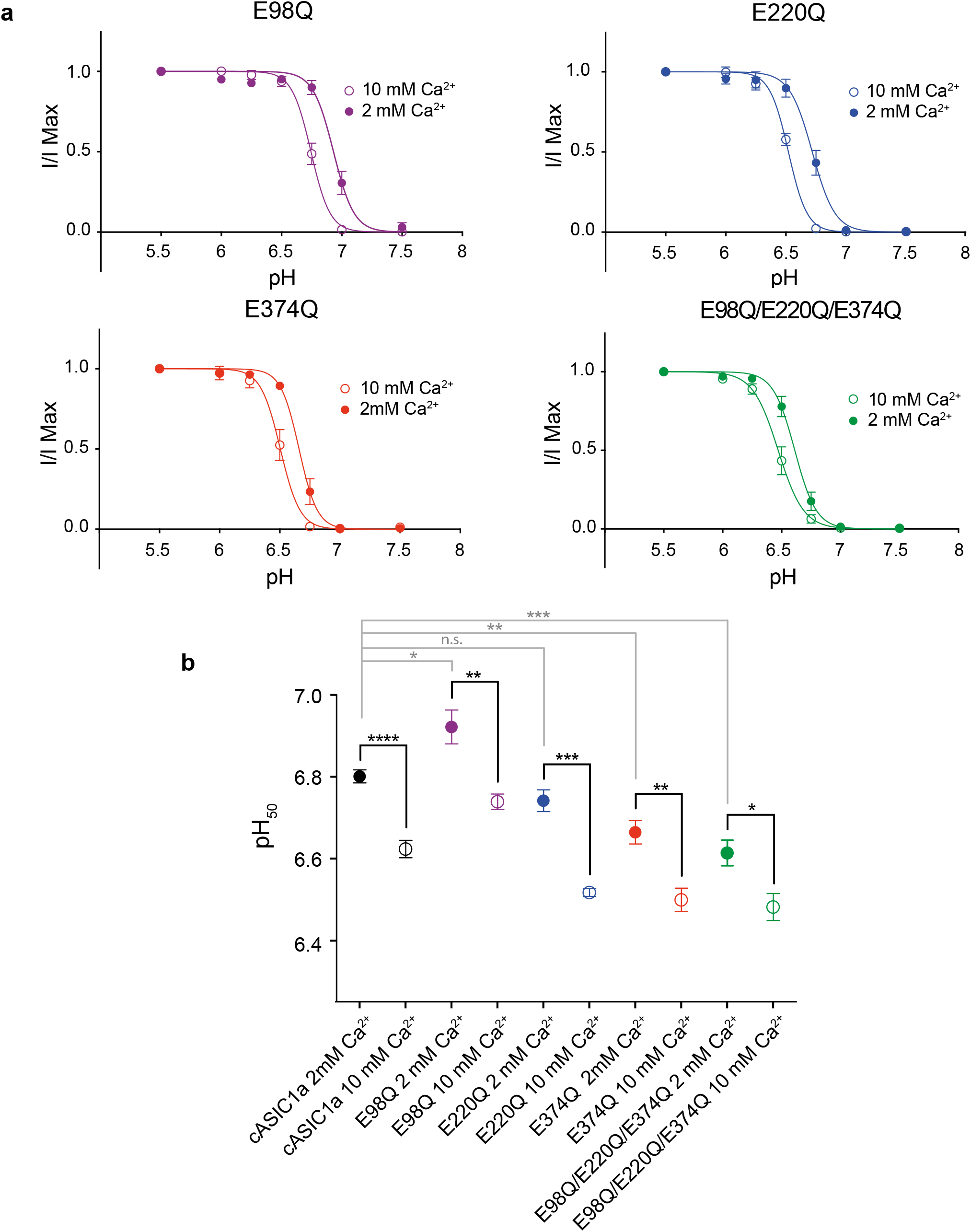
Gating modification by Ca^2+^ in cASIC1a neutralization mutants. **a,** Dose response data in 2 or 10 mM Ca^2+^ for cASIC1a mutant channels. Data were collected in whole-cell patch clamp configuration from adherent CHO-K1 cells transfected with cDNA for mutant channels. **b,** Unpaired t-test results (two-sided) comparing changes in pH50 for cASIC1a (dose response data shown in Figure 1A) and mutant channels in 2 vs 10 mM Ca^2+^ (black lines) or comparing pH50 of cASIC1a to individual mutant channels in 2 mM Ca^2+^ (grey lines). For all experiments, error bars represent SEM, n = 4-8 cells. For comparisons of pH50 values for each channel at 2 or 10 mM Ca^2+^, p values and 95% confidence intervals in parentheses are p < 0.0001 (−0.237 to −0.118) for cASIC1a, 0.0078 (−0.296 to −0.070) for E98Q, 0.0002 (−0.293 to −0.156) for E220Q, 0.0022 (−0.255 to −0.075) for E374Q), and 0.0165 (−0.234 to −0.030) for E98Q/E220Q/E374Q. For comparisons of pH50 values between channels at 2 mM Ca^2+^, p values and 95% confidence intervals in parentheses are p < 0.0405 (0.008 to 0.234) for cASIC1a and E98Q, 0.0935 (−0.132 to 0.013) for cASIC1a and E220Q, 0.0036 (−0.213 to −0.060) for cASIC1a and E374Q, and 0.0006 (−0.268 to −0.107) for cASIC1a and E98Q/E220Q/E374Q.

Our electrophysiological results highlight the complex relationship between divalent cation binding and gating in ASICs. We suggest that evidence of divalent cation binding within regions of the channel that undergo pH-dependent conformational changes as well as differences in Ba^2+^ sites observed between resting and desensitized channels demonstrate that ion occupancy is at least partly state-dependent and indicative of an overlap between pH-dependent gating and ion coordination. Furthermore, for E98Q, E374Q and triple mutant channels, changes in pH50 values in 2 mM Ca^2+^ as well as the less significant effect of Ca^2+^ on gating indicate that these residues both participate in proton-dependent gating and may contribute to modulation of gating by extracellular Ca^2+^. However, our electrophysiological results fall short of conclusively identifying residues within the acidic pocket or central vestibule that are essential to the mechanism by which divalent cations modulate channel gating. As such, additional experiments are required to generate a comprehensive understanding of the molecular mechanisms by which divalent cations modulate ASICs.

### State-dependent chloride binding

Cl^−^ is the predominant anion in the CNS, with extracellular concentrations maintained at ~ 120-130 mM(33, 34). Additionally, Cl^−^has been shown to modulate the function and structural stability of kainite receptors via a binding site at a subunit interface(34). In structures of ASIC channels in open and desensitized states, a bound Cl^−^ ion is located within the thumb domain coordinated by Arg 310 and Glu 314 of helix α4 and by a neighboring subunit via Lys 212(4, 18, 19) (Figure 5A-B). Disrupting Cl^−^ binding at this site alters pH-dependent gating, speeds desensitization and attenuates channel tachyphylasis(17). Notably, no electron density for Cl^−^ was observed in structures of a resting ASIC1a channel at high pH(23). Additionally, the rearrangement of thumb helices α4 and α5 to form the resting channel’s expanded acidic pocket at high pH disrupts the subunit-subunit interface at the carboxy-terminus of α5 and alters the architecture of the canonical and highly conserved Cl^−^ binding site (Figure 5B). These results suggest that Cl^−^ binding within the thumb domain of ASICs may be state-dependent and demonstrate how Cl^−^ may play a role in channel gating by stabilizing the collapsed conformation of the acidic pocket at low pH.

**Figure 5.**
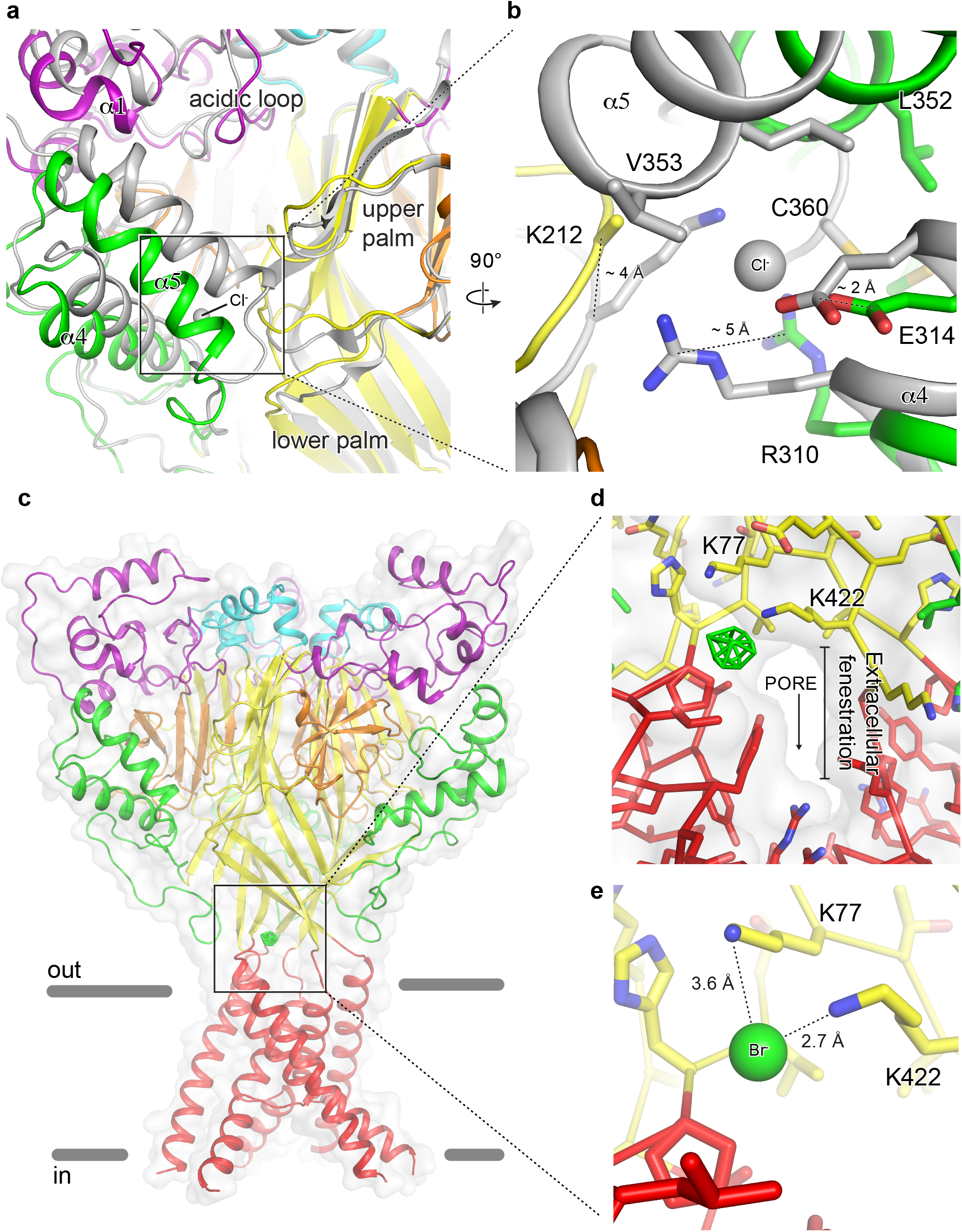
Anion binding site of a resting chicken ASIC1a channel. **a-b,** The acidic pocket (**a**) and canonical thumb domain Cl^−^ binding site (**b**) of superposed resting and desensitized (grey) channels. **c-e,** Br^−^ anomalous difference peak (green mesh) contoured at 5 σ and mapped on a resting channel at high pH (**c**) and detail views of the Br^−^ binding site with (**d**) and without (e) surface representation. One anion site, of the three, is shown.

To further probe Cl^−^ binding to the resting channel at high pH, we soaked Δ25 crystals in 150 mM NaBr. Consistent with 2Fo-Fc density maps from Δ25-Ba^2+^ and Δ25-Ca^2+^ x-ray structures(23), anomalous difference maps subjected to real space threefold averaging did not indicate the presence of a bound Cl^−^ ion at the canonical thumb domain site. Surprisingly, further inspection of these difference maps revealed strong peaks located at the mouth of the extracellular fenestrations, just above the TMD (Figure 5C). Placement of a Cl^−^ ion at this high pH binding site off of Lys 77 and Lys 422 partially occludes the narrow extracellular fenestrations that provide access for Na+ ions to the channel pore (Figure 5D-E) and that undergo a dramatic expansion upon channel activation(23). Furthermore, a recent study demonstrated that Lys 422, positioned less than 3 Å from the high pH Cl^−^ site, is important for inhibition of ASIC1a by ibuprofen(35), indicating that this region of the channel is critical for proton-dependent gating and serves as a binding site for channel modulators. Moreover, while the biological significance of a high pH Cl^−^ site on the resting channel remains unknown, the absence of a bound Cl^−^ at the canonical low pH site in crystal structures of the resting channel at high pH demonstrates how Cl^−^ ions, in addition to protons and divalent cations, play an important role in the pH-dependent function of ASICs.

## Discussion

Here we map binding sites for anions and divalent cations, determined by anomalous scattering x-ray crystallography, on x-ray structures of resting and desensitized ASIC1a channels at high and low pH, respectively. At high pH, each subunit of the resting channel harbors three binding sites for divalent cations, two of which are within the expanded conformation of the acidic pocket, and one is within the central vestibule. At low pH, the desensitized channel contains a single Ba^2+^ site at the edge of the acidic pocket. Thus, upon transition from the high pH resting state to the low pH desensitized state the divalent cation binding site in the central vestibule is lost and the there is a reduction in the number of binding sites in the acidic pocket, consistent with the notion that divalent cations bind to and stabilize the high pH resting state of the channel.

We also demonstrate that the high pH resting state of the channel lacks a bound Cl^−^ ion within the thumb domain(4). This binding site for Cl^−^, or Br^−^, is occupied in low pH desensitized and MitTx-bound open states, the latter of which was determined at pH 5.5. Electrophysiological studies have shown that this Cl^−^ site is important for ion channel desensitization kinetics and tachyphylaxis(17). When we carried out anomalous scattering x-ray crystallography experiments to study the occupancy of the thumb domain Cl^−^ site at high pH we found no evidence of ion binding, demonstrating that the thumb domain Cl^−^ site is unoccupied at high pH. The absence of a bound Cl^−^ within the thumb domain at high pH is likely due to conformational changes at the Cl^−^ site resulting from the pH-dependent expansion of the acidic pocket. Fortuitously, our crystallographic experiments uncovered a halide binding site at the extracellular fenestrations of the ion channel pore in the high pH structure of the resting channel. Further experiments are required to determine the significance of these halide binding sites on ion channel function.

A recent report demonstrated a binding site for Ca^2+^ within the pore of rat ASIC3 located within a ring of Glu residues that correspond to Gly in cASIC1a(16). These results further corroborate the observation of a Ca^2+^-dependent block reported previously for ASIC3 channels(15). Intriguingly, neither our anomalous diffraction data nor inspection of electron density maps from the Δ25-Ca^2+^ x-ray structure (pdb 5WKV)(23) indicate the presence of divalent cations near or within the channel pore as has been previously proposed(13). However, lacking structural information from an open ASIC1a channel in complex with divalent cations, we are unable to rule out the possibility of a binding site at the pore unique to the open channel.

Finally, while both Ba^2+^ and Ca^2+^ modify ASIC gating and have similar coordination requirements, Ba^2+^ has a slightly larger ionic radii(36) and it is possible that Ba^2+^-based anomalous diffraction experiments may preclude the detection of all Ca^2+^ binding sites. Therefore, while anomalous difference peaks confirm the presence of Ba^2+^ sites in resting and desensitized x-ray structures, we acknowledge the possibility of Ca^2+^ sites on ASIC1a that are incompatible with Ba^2+^ binding and thus undetected by our current experimental methods.

State-dependence for both anion and cation binding suggests an interplay between ion binding and proton-dependent gating in ASIC1a. Moreover, our results show that binding sites for divalent cations are positioned within electrostatically negative regions of the channel that undergo substantial conformational changes during gating(23) highlighting potential mechanisms of modulation. Therefore, while our results fall short of providing a detailed molecular mechanism or identifying individual residues responsible for modulation of ASICs by endogenous ions, these data underscore the complexity of the mechanisms underlying ionic modulation of ASICs and highlight the importance of additional experimentation to further improve our understanding of the relationship between ion binding and channel gating.

**Supporting information figure 1:** 5WKX validation report.

**Supporting information figure 2:** 5WKY validation report.

**Supporting information figure 3:** 6CMC validation report.

## Acknowledgements

We thank A. Goehring, D. Claxton and I. Baconguis for initial construct screening and advice through all aspects of the project, L. Vaskalis for help with figures, H. Owen for manuscript preparation and all Gouaux lab members for their support. We thank all reviewers for their time as well as for their comments and suggestions. We acknowledge the Berkeley Center for Structural Biology and the Northeastern Collaborative Access Team for help with x-ray data collection. E.G. is an investigator with the Howard Hughes Medical Institute.

